# CMash: fast, multi-resolution estimation of k-mer-based Jaccard and containment indices

**DOI:** 10.1101/2021.12.06.471436

**Authors:** Shaopeng Liu, David Koslicki

## Abstract

*K*-mer based methods are used ubiquitously in the field of computational biology. However, determining the optimal value of *k* for a specific application often remains heuristic. Simply reconstructing a new *k*-mer set with another *k*-mer size is computationally expensive, especially in metagenomic analysis where data sets are large. Here, we introduce a hashing-based technique that leverages a kind of bottom-*m* sketch as well as a *k*-mer ternary search tree (KTST) to obtain *k*-mer based similarity estimates for a range of *k* values. By truncating *k*-mers stored in a pre-built KTST with a large *k* = *k*_*max*_ value, we can simultaneously obtain *k*-mer based estimates for all *k* values up to *k*_*max*_. This truncation approach circumvents the reconstruction of new *k*-mer sets when changing *k* values, making analysis more time and space-efficient. For example, we show that when using a KTST to estimate the containment index between a RefSeq-based microbial reference database and simulated metagenome data for 10 values of *k*, the running time is close to 10x faster compared to a classic MinHash approach while using less than one-fifth the space to store the data structure. A python implementation of this method, CMash, is available at https://github.com/dkoslicki/CMash. The reproduction of all experiments presented herein can be accessed via https://github.com/KoslickiLab/CMASH-reproducibles.

## 1 Introduction

*K*-mers, contiguous strings of DNA or RNA of length *k*, are frequently utilized in computational biology for a variety of purposes including in genome assembly [12, 17, 18], metagenomic sequences classification [9, 19, 26], motif discovery [11, 27], and large-scale genomic comparisons [20, 25]. A number of hashing-based techniques such as MinHash [5], Bloom filter [4], and Count-Min Sketch [8] have been developed or adopted for efficient computation of *k*-mer based similarity methods. In each such application, the first step is to collect a set of *k*-mers from input sequences. Importantly, it has been found that algorithm performance depends critically on the choice of size *k*. Indeed, various heuristic and empirical strategies have been introduced to find optimal *k*-mer sizes that increase performance in certain application areas [7,24,28]. However, whenever a new *k* size is selected, each computational technique requires reconstructing the *k*-mer based data structure and rerunning the analytical pipeline, leading to computational inefficiencies.

In particular, hashing-based *k*-mer methods that compute measures of similarity of genomic and metagenomic data (such as the Jaccard and containment indices) have been demonstrated to extract valuable insight from metagenomic data [3, 19, 21]. Multiple hashing based techniques involving the estimation of Jaccard index and/or other *k*-mer derivatives have been developed. For example, Mash [20], Sourmash [21], and Skmer [23]. Several efforts have been made to improve the efficiency for single *k* value hashing method, such as b-bit wise MinHash [16] and Dashing with HyperLogLog sketches [1]. In these cases too, however, each time a new *k* size is utilized, the entire computational processes needs to be repeated.

To address this problem, we combine a modified MinHash technique (ArgMinHash) and a data structure called a *k*-mer ternary search tree (KTST), which allows Jaccard and containment indices to be computed at multiple *k*-mer sizes efficiently and simultaneously. In Fig. 1 we provide a high-level description of how we accomplish this: first, we randomly subsample *k*-mers based on a large *k* size *k_max_* (Fig. 1b) to build *k*-mer sketches. The sketch elements (i.e. *k_max_*-mers) are then inserted into a KTST (Fig. 1c), which allows for efficient prefix lookups. A prefix lookup in the KTST effectively truncates a *k_max_*-mer resulting in a smaller *k*-mer (Fig. 1d). This allows us to efficiently compute *k*-mer sketches for every *k* ≤ *k_max_* (Fig. 1e). Combined with the containment MinHash approach [14], we can estimate the Jaccard and containment indices for all *k* < *k_max_* without requiring explicit re-computation of each single *k* value.

**Fig. 1.**
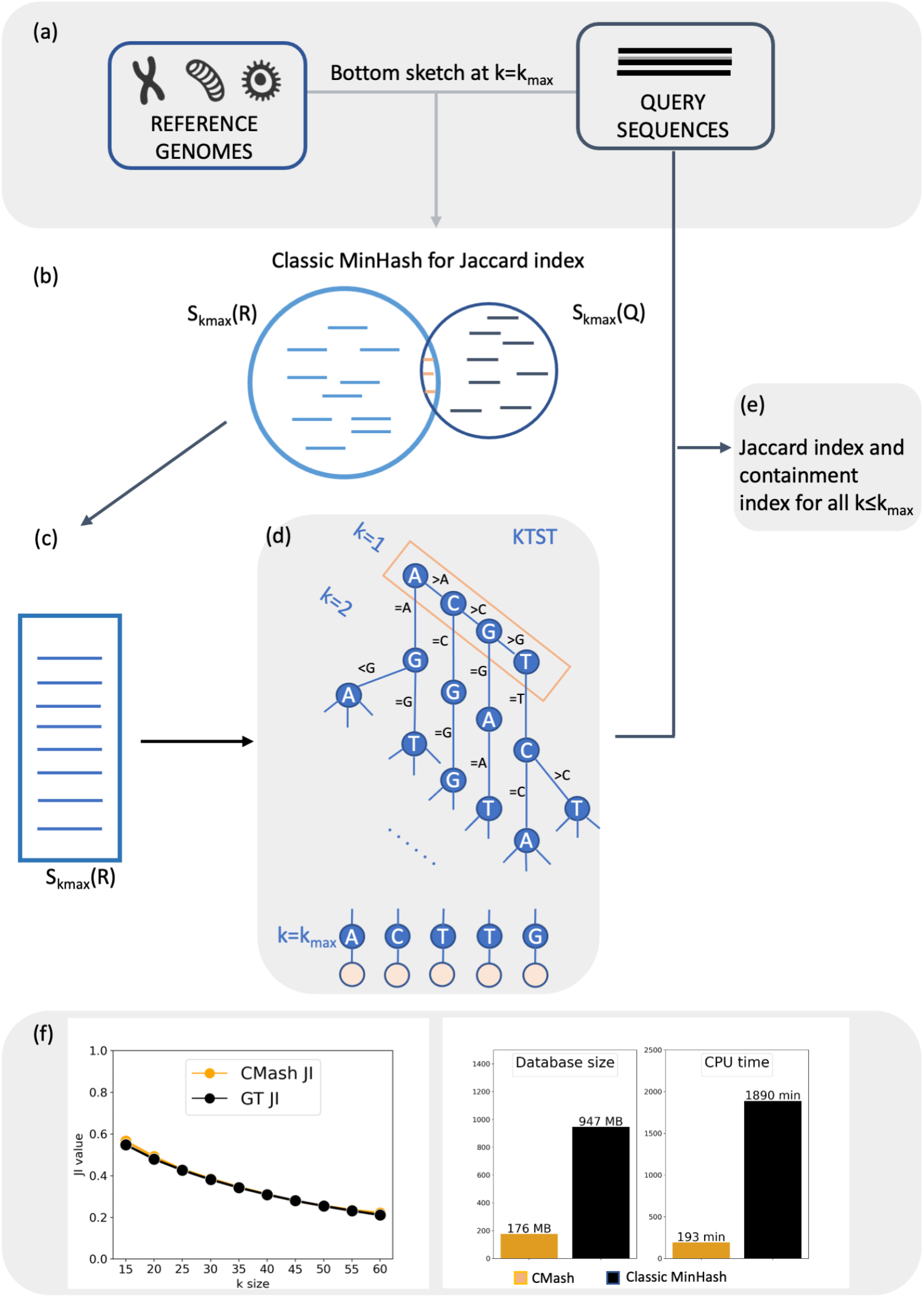
Overview of the CMash algorithm. (a) The input to CMash are genomes or sequencing reads. (b) Random samples of *k*-mers using a modified bottom *n* sketch can also be used for the classic MinHash algorithm. (c) For some large *k* value *k*_*max*_, one and only one *k*-mer sketch of the reference data will be constructed and inserted into a KTST. (d) All *k*-mer sketches corresponding to a smaller *k* value will be obtained by a prefix lookup in the KTST. (e) For *k* < *k*_*max*_, *k*-mers from the query data are streamed through the KTST resulting in (f) reliable estimates for a range of *k*-mer sizes with greater computational efficiency.

This truncation-based method turns out to be a biased estimator of the *k*-mer based similarities. However, in our empirical analysis, we find that the CMash estimate of the Jaccard and containment index does not deviate significantly from the ground truth, indicating that this approach can give fast and reliable results with minimal bias. Compared to our previous MinHash-based approximation to the containment index [14], we find that the CMash estimate for ten *k* values is approximately ten times faster and requires only one-fifth of space to store the reference database.

Importantly, this approach can be generalized to more than similarity computation: many sketching, *k*-mer, or shingling-based approach may adopt our method to avoid the need to re-compute *k*-mer sets when changing the *k* size. As such, this probabilistic data analysis approach should find application outside of metagenomics (the application we focus on) to genomics more broadly, as well as other applications that utilize a *k*-mer or shingling approach.

In summary, we demonstrate how this CMash technique can be applied to several widely utilized tools (e.g. Mash Screen [19], Sourmash [21]) and will help to speed up *k*-mer based computation when multiple *k* sizes are needed. A proof of concept implementation of the algorithm and data structure is freely available at https://github.com/dkoslicki/CMash.

## 2 Method

Here we describe our algorithmic approach, but first we recall a few necessary definitions.

### 2.1 Preliminaries

#### 2.1.1 Jaccard and containment index

In computational biology, *k*-mers are consecutive substrings of length *k* of nucleotides 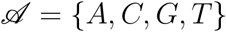. The similarity between genomic data can be measured by the similarity of their respective *k*-mer sets: the collection of all distinct *k*-mers appearing as contiguous substrings in the data. If *A* is a collection of strings on the alphabet 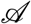, then *A^k^* is defined to be the set of all unique *k*-mers in *A*. In this entire section, most of the definitions given apply to arbitrary sets, but with the genomic application area in mind, we often suppress the superscript and write *A* instead of *A^k^* for simplicity, with the implicit understanding that a set of *k*-mers depends on the *k* value chosen.

The Jaccard index (JI) measures the similarity of two sets by comparing the relative size of the intersection over the union [5]. For two non-empty finite sets *A* and *B*, the Jaccard index is defined as 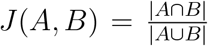. Hence, 0 ≤ *J*(*A, B*) ≤ 1 with larger values indicating more overlap. Similarly, the containment index (CI) of *A* in *B* (with *A* non-empty) measures the relative size of the intersection over the size of 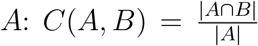. So 0 ≤ *C*(*A, B*) ≤ 1 with larger values indicating that more content of *A* resides in *B*. If the cardinality of both *A* and *B* are known, the Jaccard index and containment index are interchangeable:

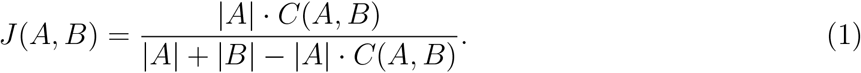

When applied to sets of *k*-mers, we call out the dependence on *k* with the following definitions:

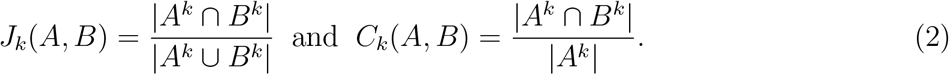

#### 2.1.2 Classic MinHash algorithm for the Jaccard index

For very large sets *A* and *B* (such as *k*-mer sets for moderate to large *k* derived from genomic data), computing the Jaccard index directly can be computationally taxing. To circumvent this, Broder proposed MinHash to efficiently estimate the Jaccard index for large sets [5]. MinHash uses a random sampling process: first, we fix a constant 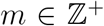 (*m* is usually called sketch size) and select a family of *m* min-wise independent hash functions 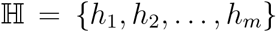 whose domains contain |*A* ∪ *B*|. Then, we define the MinHash sketch of a set *A* as the element (ties can be solved by lexicographic order) in *A* that cause some *h_i_* to have the minimum value on *A*. More formally, define 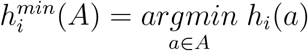. Next, define *m* random variables 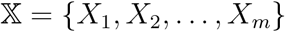, such that:

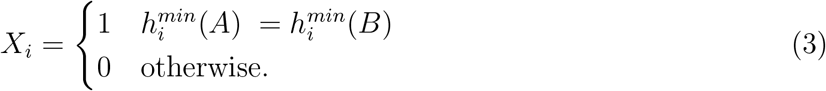

The probability of a MinHash collision (i.e. 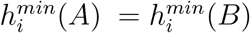) is an unbiased estimate of *J* (*A, B*):

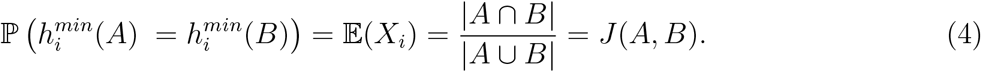

In practice, a “bottom sketch” strategy, originally proposed by Broder [5], is commonly used to implement the MinHash algorithm. Instead of using *m* hash functions, all *k*-mers from a given set *A* are passed through a single hash function and the smallest *m* hash values (instead of elements) are stored in a sorted sketch *S_b_*(*A*) of size *m*. The probability that sketch *S_b_*(*A*), *S_b_*(*B*) share a hash value represents the probability of random sampling a shared element from the union of set *A* and *B*. So, the resemblance of set *A*, *B* can be quickly estimated by counting the matched values between *S_b_*(*A*) and *S_b_*(*B*).

This efficient approach has found use in, for example, metagenomics where hundreds of thousands of microbial genomes may under consideration. For example, both Sourmash [21] and Mash Screen [19] maintain hash *value* sketches of all input genomes for comparison. However, *k*-mer information is lost during if one only considers hash values, instead of elements leading to minimal hash values. Herein, we will show how we can benefit from using a *k*-mer sketch instead of a hash value sketch in similarity analysis.

We now define a bottom *m k*-mer sketch. Let *A^k^* be the set of all *k*-mers derived from a set of sequences/string *A* and define MIN_*m*_ (*A^k^*) as the set of the *m* elements corresponding to the *m* smallest hash values in set 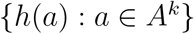. Namely, for *m* a given sketch size and *k* the *k*-mer size, the *k*-mer MinHash sketch of *A* is defined to be

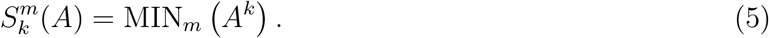

We may suppress *m* and *k* for notational simplicity.

### 2.2 Containment MinHash

Though the MinHash approach gives an unbiased estimation of the Jaccard index, its performance may degrade considerably when *A* and *B* are of significantly different sizes [14]. More robust estimation of *J*(*A, B*) can be obtained through *C*(*A, B*), the containment index of *A* in *B*. This strategy is called “containment MinHash” [14]. We detail this procedure now. Given a fixed *k*-mer size and two nonempty distinct sets of strings *A* and *B* on the alphabet 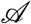 such that |*A^k^*| ≤ |*B^k^*|, we first compute 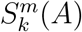, the bottom sketch of the smaller set. Next, we can stream all elements in the set *B* over 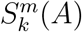 to estimate *C*(*A, B*). Since 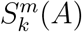 is a uniform random sample from set *A*, the proportion of elements in 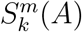 that are found in set *B* is an unbiased estimator of the containment index. Namely,

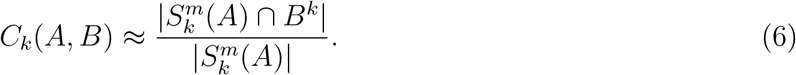

To be noted, this streaming method is an efficient algorithm for the estimation of the containment index in metagenomic settings and is utilized by Mash Screen [19], Metalign [15], etc. Finally, we can take advantage of equation (1) to compute *J_k_*(*A, B*) based on the containment index and the cardinalities of set *A* and *B* (which can be quickly approximated by fast cardinality estimation such as Hyperloglog [10]). In CMash, we use this contaiment MinHash approach for JI estimation considering its metagenomic analysis setting.

### 2.3 CMash

The approach we call CMash consists of two main components: first is the aforementioned *k*-mer MinHash sketches, and second a traditional ternary search tree applied to sets of *k*-mers.

#### 2.3.1 ArgMinHash

We now detail the first half of the CMash approach: a data structure we call “ArgMinHash” that utilizes *k*-mer MinHash sketches. In particular, there is an important but subtle difference between the aforementioned MinHash bottom *m* sketches and the *k*-mer MinHash sketches utilized by CMash. In particular, the definition in equation (5) shows that the bottom *m* sketch utilized by the containment MinHash (or even MinHash itself) are comprised of the smallest *m hash values*. In contrast, the sketches utilized by CMash are comprised of *elements* of a set that hash to small values. This difference is key to allowing a truncation based approach. Indeed, if we used a sketch comprised of hash *values* of *k*-mers, truncating these hash values would have no relationship at all to the hash values obtained from truncated *k*-mers.

More formally, let *A^k^* be the set of all *k*-mers derived from a set of sequences/strings *A*. A hash function *h* with domain containing *A^k^* induces an order on *A^k^*. If collisions are present, we can impose an additional ordering (say, lexicographic) to break ties. Then we define 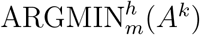 as the set of *m* smallest, according to the ordering imposed by *h*, elements of the set *A^k^*. Then for *m* a given sketch size and *k* the *k*-mer size, the “argmin-bottom” sketch of *A* is defined to be

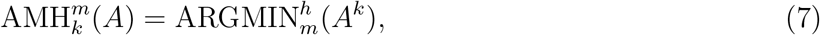

where AMH is an abbreviation for “ArgMinHash.”

#### 2.3.2 K-mer ternary search tree

Given a set of collections of sequences *D* = {*A*_1_, . . . , *A_N_*}, here thought of as genomes of *N* different (micro)organisms, we populate a single ternary search tree KTST with the sketches 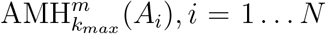 for a fixed sketch size *m* and a fixed (large) *k_max_*. Recall that a ternary search tree is a data structure that allows fast (average *O*(log *n*)) lookup of prefixes so that every root to leaf path (equivalently, node) represents a *k*-mer. Furthermore, nodes in KTST can be labeled with which elements of *D* contain the prefix defined by that node. We further associated a sequence of counters 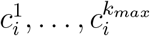 to each *A_i_* in *D*. We further accelerate prefix queries by populating a bloom filter with every *k*-mer defined by nodes in the KTST.

Note that by inserting the sketches 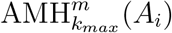 into the KTST, we have effectively computed proxies to 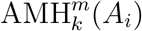 for each *k* ≤ *k_max_*. Indeed, we obtain new sketches for a smaller *k*-mer size *k* by truncating the KTST to a depth of *k* (Fig. 1d).

We can then approximate the containment index of each reference *A_i_* in some other set of sequences *B* (thinking of *B* as a large genomic data set) in the following way: The *k*-mers of *B* for each *k* = 1, 2, . . . *k_max_* are streamed through the KTST similar to the aforementioned Mash Screen (see Fig. 1e). When a *k*-mer is found to correspond to a node in the KTST, each of the counters 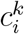 is incremented for each *A_i_* associated with that node/*k*-mer. After the streaming is complete, we will have that

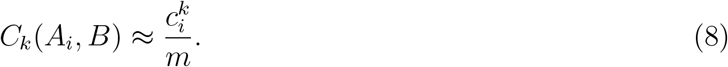

In doing so, in a single stream over the input data *B*, we are able to approximate *C_k_*(*A_i_*, *B*) for each *A_i_* and each *k* = 1, . . . , *k_max_*. If during the construction of the sketches 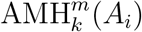, we also store the cardinality of 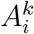, we can obtain the Jaccard indices *J_k_*(*A_i_*, *B*) as well.

The ability to estimate Jaccard or contaiment indices for multiple *k*-mer sizes (up to some maximum *k_max_*-mer size) motivates the multi-resolution nature of CMash. Indices can be calculated for both small *k*-mer sizes (low resolution), and large *k*-mer sizes (high resolution) utilizing a single data structure.

#### 2.3.3 Biased nature of the estimate

There is no reason to think that the estimate in equation (8) will be unbiased. Indeed, while 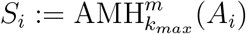 is truly a random sample of *m* elements from the set 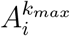 and so the estimate given in equation 8 corresponds exactly to MinHashing with *k* = *k_max_*, truncating the elements of *S_i_* to *k*-length prefixes will not be a random sample of *m* elements from 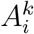 due to duplicated prefixes. Consider *A* = {AATAAG} with *k_max_* = 3 and *m* = 1: every one of the four 3-mers AAT, ATA, TAA, AAG has equal probability of being selected. As such, truncating these to 2-mers results in AA appearing with probability 50% in the truncated 3-mers. In contrast, the frequency of AA in *A*^2^ is only 25% as 4 distinct 2-mers *A*^2^ = {AA, AT, TA, AG}. Though the truncation step will inevitably introduce some bias in the estimation, the gain in speed overwhelms the small sacrifice to accuracy, which we empirically verify in the next sections.

### 2.4 Theoretical analysis of CMash

Theoretically and practically, a truncation-based estimate of the Jaccard similarity will introduce data-dependent bias. Consider an arbitrary sequence data *A* and let *A^k^* denote the set of all distinct *k*-mers of length *k* in *A*. Obviously, *A*^1^ = {*A, C, G, T*}. Similarly, *A^k+L^* denote the set of all distinct (*k* + *L*)-mers in *A*. Let (*A^k+L^*)_1…*k*_ denote the distinct *k*-mers obtained by directly truncating all elements in *A^k+L^* from length (*k* + *L*) to *k*. In an ideal situation where no two elements share the same prefix of length *k*, the truncated *k*-mer set (*A^k+L^*)_1…*k*_ is exactly *A^k^*. Unfortunately, this will not happen in most cases where duplicate prefixes will be introduced during the truncation, leading to estimation deviance. In this section, we will show how this truncation-introduced bias correlates with the the truncation length *L* as well as the input data themselves.

#### 2.4.1 Bias in truncation-based Jaccard index

First we define a prefix relationship between two *k*-mers of different lengths. For *k*-mer *M_k+L_* of length *k* + *L* and *N_k_* of length *k*, if *N_k_* is a prefix of *M_k+L_*, we can truncate *M_k+L_* by length *L* to get *N_k_*, which is written as *N_k_* = (*M_k+L_*)_1…*k*_. We may suppress the length subscript for notational simplicity. Namely: *N* = *M*_1…*k*_.

We then define right extensions: for a given *k*-mer *X* of length *k*, and 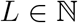, we use 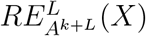 to denote all (*k* + *L*)-mers in the set *A^k+L^* that have *X* as prefix. That is to say,

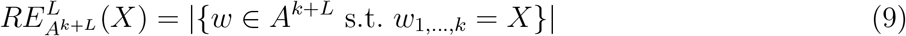

Now, we can quantitatively describe the bias in the truncation-based method. Let 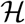 be a family of suitable hash functions (eg. min-wise independent). Given two arbitrary non-empty genome/sequence files *A* and *B* (and the *k*-mer sets *A^k+L^* and *B^k+L^* for any arbitrary positive integers *k, L*), if we truncate all *k*-mers from length *k* +*L* to *k*, the truncation-based Jaccard index truncated from *k* +*L* to *k*, denoted by *JI*(*A, B*)_*trunc*(*k*+*L*→*k*)_ can be computed in the following:

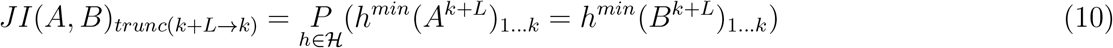

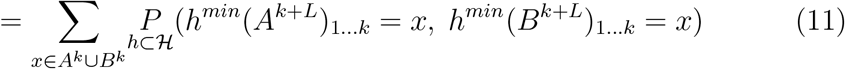

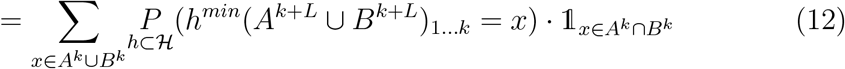

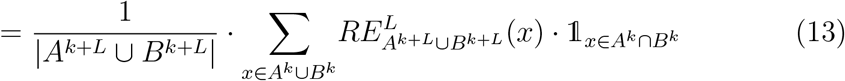

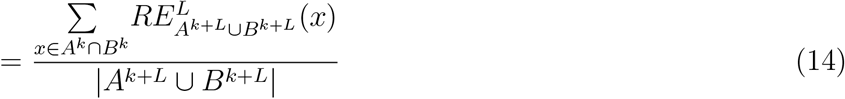

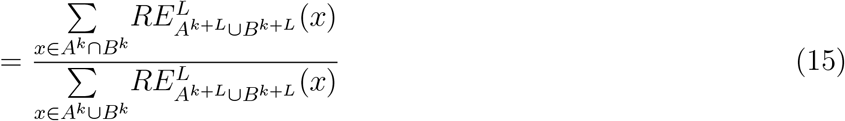

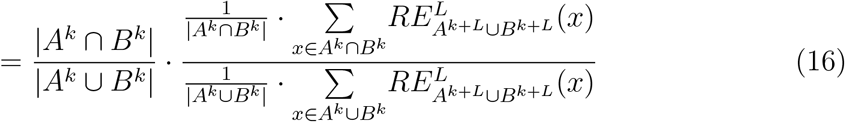

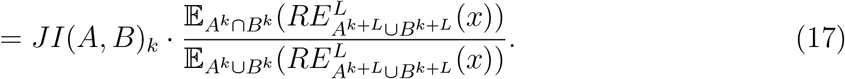

Briefly, the truncation-based estimation of Jaccard index utilized by CMash will bring a multiplicative bias factor upon the classic MinHash estimation of the Jaccard index as shown in eqn. 17. This bias factor reflects the imbalance of prefixes (i.e. truncated *k*-mers) distributions between the intersection and the union of the original *k*-mer sets (before truncation). The bias factor implies that CMash will be more reliable when there are few duplication after truncation (in this case, expected right extension of any prefix tends to be 1) or when *A* and *B* have relative high similarity (namely, the intersection well represents the union). In practice, CMash is more reliable when using large *k* values or running with closely related data. To demonstrate this, we applied this approach to 31 genomes within the genus Brucella with multiple *k* values (Fig. 2) and showed that CMash can robustly function to estimate Jaccard indices for multiple *k* values simultaneously. Considering the unavoidable variance due to random sampling in MinHash algorithm, the bias in CMash may not be an obstacle empirically.

**Fig. 2.**
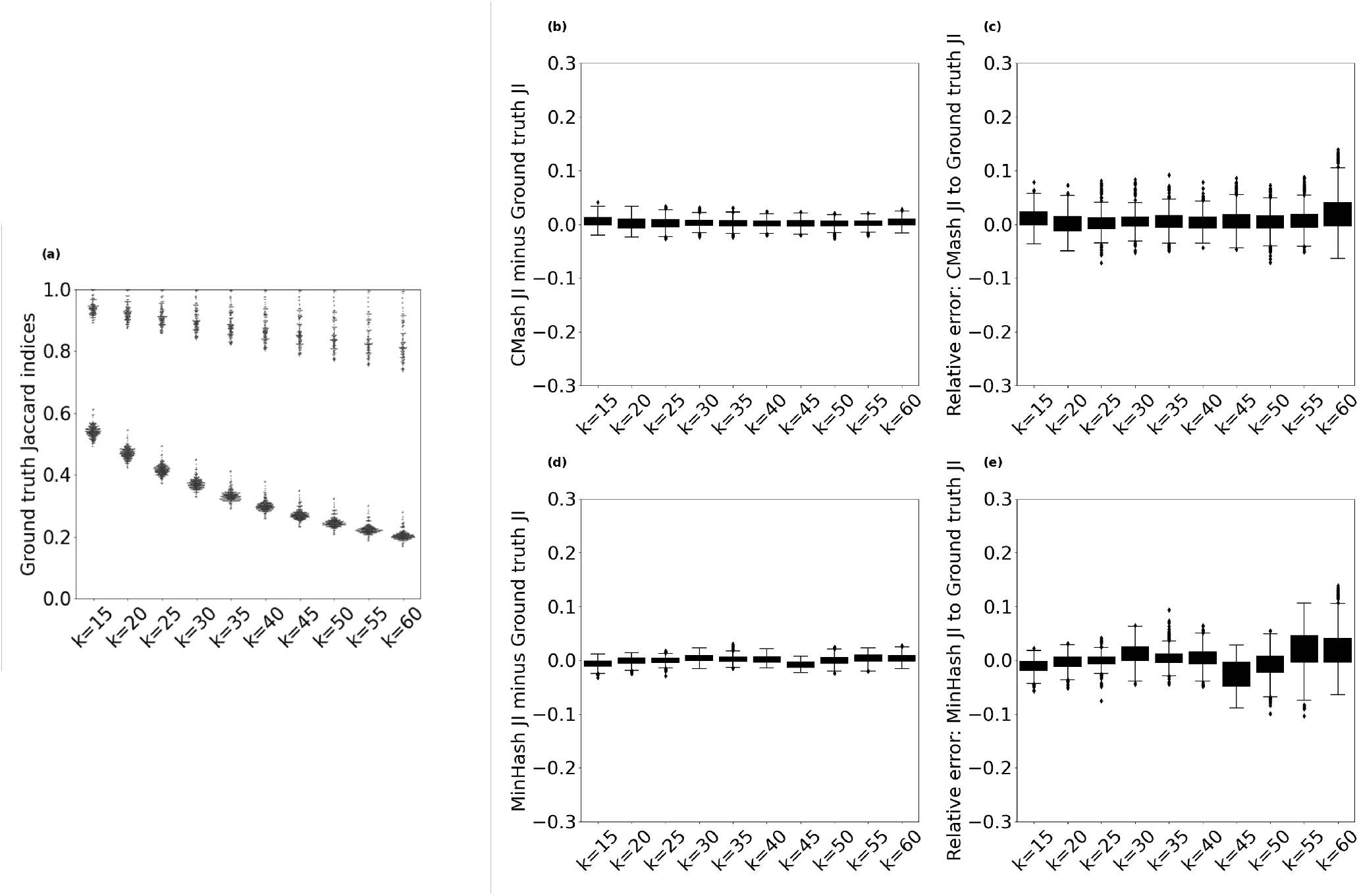
Jaccard indices obtained through CMash and MinHash compared to the ground truth on all pairs of 30 Brucella genomes. (a) The ground truth Jaccard indices as a function of *k*-mer size from *k*=15 to *k*=60. (b) Boxplot of JI value differences between CMash and the ground truth. (c) Boxplot of relative errors of CMash compared to the ground truth (d) The relative error of CMash compared to the ground truth. (d) Boxplot of JI value differences between MinHash and the ground truth. (e) Boxplot of relative errors of MinHash compared to the ground truth.

#### 2.4.2 Bias in truncation-based containment index

Similar to Mash Screen [19], when computing the containment index (CI) of *B^k^* in set *A^k^*, performance improvements can be had if the one forms only a sketch *S*(*A^k^*) of *A^k^* and then streams the elements of *B^k^* over it, looking for matches. Hence, truncation of elements of *B^k^* is not necessary. We can then connect this streaming, truncation-based estimate of the containment index to the classic MinHash algorithm directly. To that end, let *m* be the given sketch size, and compute:

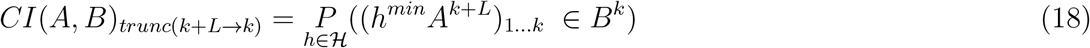

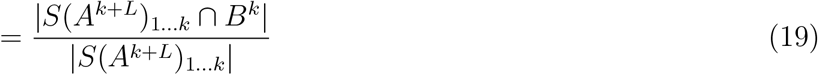

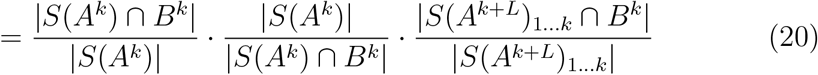

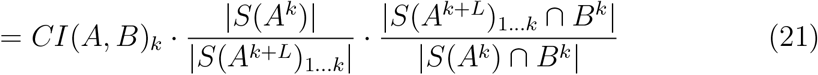

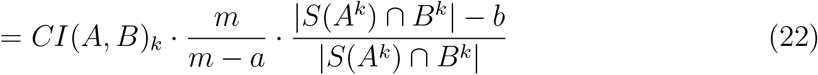

where 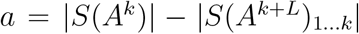 refers to the number of duplicate *k*-mers (prefixes) generated during truncating the *k*-mer sketches of *A*; and 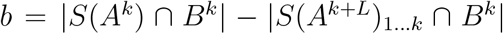 refers to the difference of cardinality of overlapping elements between the untruncated sketch and truncated sketch with *B*. Although the truncated sketch is not exact the same as the bottom sketch *S*(*A^k^*), the differences are negligible in practice due to the uniformity of the hash function(s), as we note below in section 3.

Similar to the truncated Jaccard index, the CMash estimate of the containment index will lead to a data-dependent bias factor that relies on the original *k*-mer length, the truncation length as well as the *k*-mer distribution in the input data themselves. The bias factor can be minimized when there are few duplicate prefixes (i.e. using large *k* values). Besides, a larger sketch size *m* can overwhelm the value of *a* and *b*, making the bias negligible in practice. The performance of CMash on truncated CI is examined in Fig. 3a, showing reliable estimation while being more efficient in metagenomic settings.

**Fig. 3.**
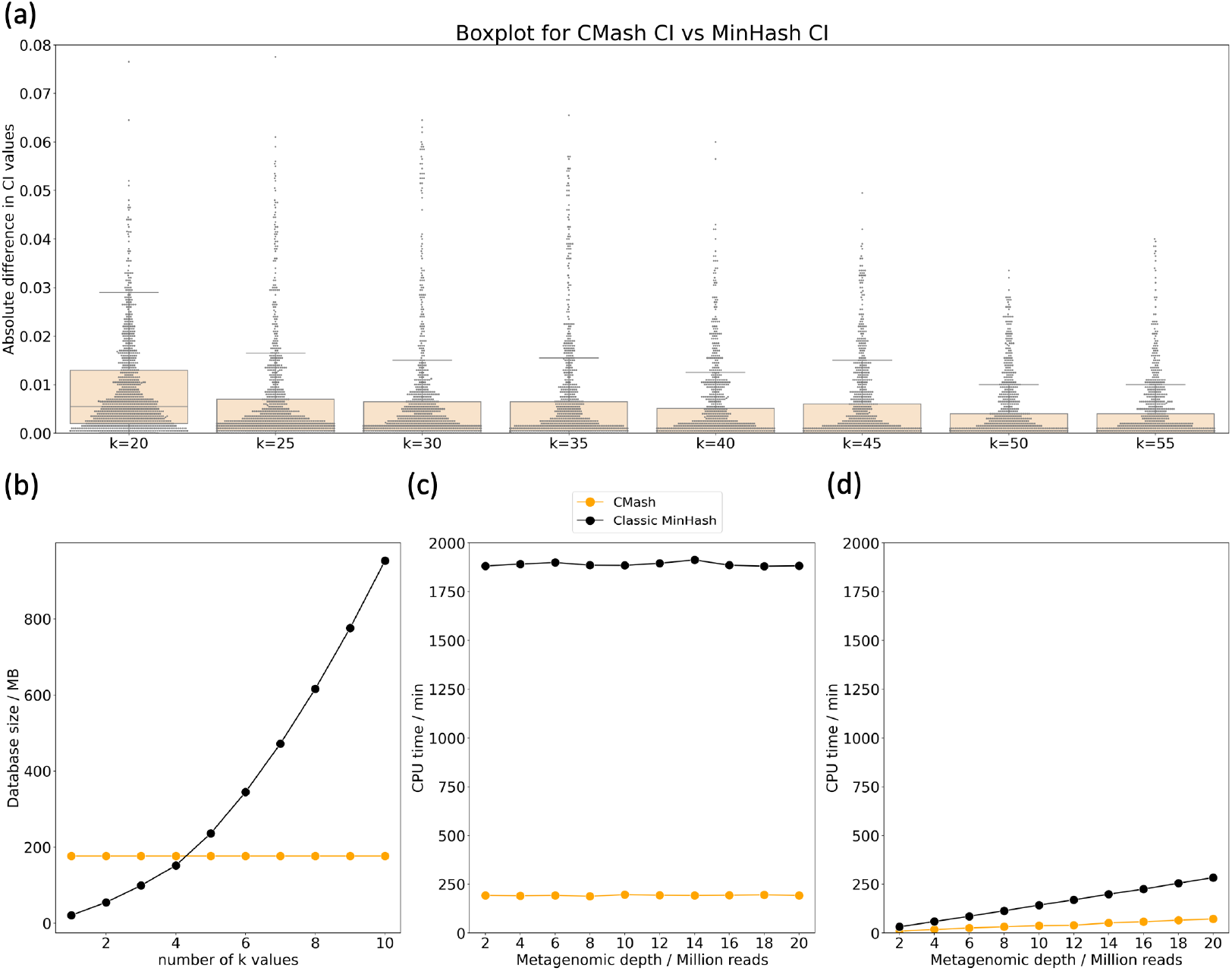
Comparison of CMash with the classic MinHash approach to quantify containment indices, along with *k* size, database creation time, and query time. The metagenomic data was simulated from 200 randomly selected genomes; and then 1000 random genomes (including the 200 true members) were analyzed for the containment index for *k* values ranging from 20 to 60. (a) Boxplot for absolute difference of CI value between CMash (*k*_*max*_=60) and the classic MinHash algorithm under different *k* values. The x-axis stands for different *k* values and y-axis stands for the absolute difference in CI. The majority of them are below 0.02. (b) Space usage for the two methods. (c) Time (per CPU minute) needed by the two methods for data structure construction. (d) Query time (per CPU minute) needed by the two methods.

## 3 Results

Here, we compare the results from CMash using the truncation-based method to both the classic MinHash estimation as well as the ground truth (brute force) calculation on real and simulated data.

### 3.1 CMash accurately estimates the Jaccard index

Considering the metagenomic setting where researchers are less interested in distal relationships, we benchmarked the efficacy of CMash on a collection of organisms all belonging to the same genus: we selected the genus Brucella. A total of 31 complete or scaffold genomes were found in the NCBI GenBank database [2] and all were downloaded, except for a single genome belonging to the species *Brucella intermedia* which was discarded due to its large evolutionary distance to the remaining 30 Brucella genomes. To assess the ability of CMash to estimate the Jaccard index, we computed all pairwise Jaccard indices for this set of genomes and compared them to the true Jaccard indices which were computed in a brute force fashion. Figure 2 contains the results where the *k*-mer size ranged from 15 to 60 in steps of 5, for a *k_max_* value of *k_max_* = 60. The sketch size for CMash was *m* = 2000 by default (estimation variance decreases exponentially with increased sketch size while the computation becomes less time-/space-efficient [14]). We use canonical *k*-mers throughout: the lexicographic minimum of the *k*-mer and its reverse complement.

As expected, Fig. 2a shows that the Jaccard index between pairs of genomes decreases as *k* increases. Indeed, for a *k*-mer size equal to an input genome’s length, the Jaccard index at this *k*-mer size is equivalent to exact string matching. As an aside, the rate of decrease of the Jaccard index as a function of *k* recapitulates the evolutionary relatedness first observed in [13], which motivated the investigation contained in this manuscript.

The differences between the CMash estimate when compared to the ground truth are explained by two components: the variance introduced by the sampling-based approach of the ArgMinHash component of CMash, and the bias introduced by truncating the KTST. As seen in, Fig. 2b and c, neither of these biases are significant when estimating the Jaccard index with CMash when comparing pairs of genomes with medium or high Jaccard similarity. The higher relative error in Fig. 2c for large *k* size is due to the decrease in the ground truth Jaccard index values as shown in Fig. 2a. CMash and the MinHash estimate exactly agree at *k* = *k_max_*, so the performance characteristics are already well studied in this setting, eg. [14]. Indeed, Fig. 2b shows that the absolute differences are tightly distributed around zero. Fig. 2d and e depicts the performance of the classic MinHash method for comparison. The lower variance of CMash estimation is achieved through the containment MinHash method in section 2.2 [14].

### 3.2 CMash is significantly more efficient than MinHash

The large size of microbial genome reference databases is a constraint in metagenomic analyses due to database size directly impacting computational time. This is especially a concern when multiple *k* values are required [13, 21, 22].

To examine the ability of CMash to ameliorate these concerns with large reference databases, we analyzed simulated metagenomic reads for the containment estimation of selected reference genomes. Among all species with complete or scaffold genomes in the NCBI GenBank database [2], we selected 1000 of them spanning 26 phyla, 174 families, and 313 genera to serve as a reference database. Next, 200 of these 1000 genomes were used to simulate metagenomic samples. We used BBTools `randomreads.sh` [6] with the default metagenomic setting to simulate these datasets. In total, ten metagenomic data sets with depths ranging from 2 million reads to 20 million reads were simulated and then processed by CMash. We compared the CMash truncation-based estimate to the classic MinHash algorithm in a direct comparison: both algorithms were coded in the same programming language and using the same hash function. The choice of *k* values was slightly different than before: *k* values ranging from 20 to 60 in steps of 5 were used and *k*=15 was excluded because the probability of sharing a 15-mer merely by chance is not negligible in the metagenomic setting where the genome pool tends to be large.

We used the containment index (CI) in this experiment due to the very different sizes of the input data: it has been established that hashing approaches more accurately estimate the containment index in this such situations [14], though recall that Jaccard and containment indices can be computed from each other when the cardinalities are known (see equation (1)). Considering that the major interests in metagenomic anlaysis are for microbes which show up in the sample (usually with moderate or high CI values) and *k*-mer matches from related or random genomes is unavoidable, we compared absolute difference of CI values between CMash and the classic MinHash algorithm for all the 1000 reference genomes. While the performances from different depths are similar, we only present the results for the depth of 10 million reads. In fig. 3a, most of the absolute CI difference falls below 0.02, suggesting that CMash consistently agrees with the classic MinHash algorithm for all *k* values considered.

Given the comparable performance, the CMash results were obtained more efficiently in terms of space and running time. CMash requires only one reference database for the estimation of all *k* ≤ *k_max_* while the classic MinHash requires space linear in the number of *k* values (Fig. 3b). While CMash is currently a prototype model, the classic MinHash method is not implemented in the most memory efficient way (both hash values and *k*-mers are stored). Though the cost of the classic MinHash method was overestimated, the superiority of CMash in dealing with multiple *k* values is significant. In this experiment that used 10 *k* values, CMash used a total of 176MB in space for the reference database while the classic MinHash approach used 947MB to store all of its sketches. In addition, due to not needing to reconstruct new sketches for new *k* values, we observe in Fig. 3c that the time needed for reference construction by CMash was almost one-tenth compared to the classic MinHash. The estimation portion, depicted in Fig. 3d, was negligible in comparison to the database construction time, but here too we found CMash to be more efficient than the MinHash approach.

## 4 Conclusion

In this manuscript, we introduced CMash: an algorithm and data structure that can provide efficient multi-resolution estimation of *k*-mer based Jaccard and containment indices. It combines a bottom “argmin sketch” strategy and a prefix lookup in a *k*-mer ternary search tree (KTST) to avoid the reconstruction of the entire reference databases each time the *k*-mer size is changed. One advantage of using a KTST not explored here is that a KTST can be represented as an on-disk database, thus freeing memory for other purposes. Indeed, the minimum memory needed is the size of one leaf node in the pre-built KTST. If needed, the amount of memory utilized by a KTST can be adjusted for a trade-off between speed and memory usage.

We showed that this truncation-based method can provide results that well-approximate the ground truth in a more computationally efficient manner. We used CMash to analyze real microbial data and simulated metagenomic data and found it to give consistent and reliable estimates. While not an unbiased estimate for *k*-mer sizes smaller than the input maximum *k_max_* value, we observed that the introduced bias was negligible for genomes with moderate and high Jaccard indices.

The required space used by this approach is constant when we fix the choice of *k_max_*, regardless of the number of different *k* values that we are interested to explore. In contrast, a classic MinHash method requires space that is linear to the number of *k* values used. Similarly, the time to construct reference database is significantly improved compared to MinHash as CMash only needs to proceed the data once; hence the total running time are effectively linearly reduced with respect to the number of *k*-mer sizes when compared to MinHash. This feature is extremely helpful in metagenomic analysis where the reference database can be as large as hundreds gigabytes and the querying cost can be overwhelmed by reference construction cost.

With further theoretical analysis of the bias factor, we believe this method will be useful in many metagenomic analyses where multi-resolution estimates can illuminate evolutionary relationships. Beyond metagenomics, *k*-mer (or shingling)-based methods are utilized extensively in computer and data science, so the CMash approach should find application beyond computational biology by essentially allowing multi-resolution (in terms of *k*-mer/shingling size) queries with little sacrifice to accuracy but greatly improved efficiency.

## Notes

* This material is based upon work supported by the National Science Foundation under Grant #1664803.

### Competing Interest Statement

The authors have declared no competing interest.

### Summary of Updates

Update Fig1 for ternary search tree illustration; Update Fig2 for the way data are presented; No factual chances.

## References

1. Baker, D.N., Langmead, B.: Dashing: fast and accurate genomic distances with hyperloglog. Genome biology 20(1), 1–12 (2019)

2. Benson, D.A., Cavanaugh, M., Clark, K., Karsch-Mizrachi, I., Ostell, J., Pruitt, K.D., Sayers, E.W.: Genbank. Nucleic acids research 46(D1), D41–D47 (2018)

3. Besta, M., Kanakagiri, R., Mustafa, H., Karasikov, M., Rätsch, G., Hoefler, T., Solomonik, E.: Communication-efficient jaccard similarity for high-performance distributed genome comparisons. In: 2020 IEEE International Parallel and Distributed Processing Symposium (IPDPS). pp. 1122–1132. IEEE (2020)

4. Bloom, B.H.: Space/time trade-offs in hash coding with allowable errors. Communications of the ACM 13(7), 422–426 (1970)

5. Broder, A.Z.: On the resemblance and containment of documents. In: Proceedings. Compression and Complexity of SEQUENCES 1997 (Cat. No. 97TB100171). pp. 21–29. IEEE (1997)

6. Bushnell, B.: Bbtools. BBMap (2018)

7. Chikhi, R., Medvedev, P.: Informed and automated k-mer size selection for genome assembly. Bioinformatics 30(1), 31–37 (2014)

8. Cormode, G., Muthukrishnan, S.: An improved data stream summary: the count-min sketch and its applications. Journal of Algorithms 55(1), 58–75 (2005)

9. Dilthey, A.T., Jain, C., Koren, S., Phillippy, A.M.: Strain-level metagenomic assignment and compositional estimation for long reads with metamaps. Nature communications 10(1), 1–12 (2019)

10. Flajolet, P., Fusy, É., Gandouet, O., Meunier, F.: Hyperloglog: the analysis of a near-optimal cardinality estimation algorithm. In: Discrete Mathematics and Theoretical Computer Science. pp. 137–156. Discrete Mathematics and Theoretical Computer Science (2007)

11. Fletez-Brant, C., Lee, D., McCallion, A.S., Beer, M.A.: kmer-svm: a web server for identifying predictive regulatory sequence features in genomic data sets. Nucleic acids research 41(W1), W544–W556 (2013)

12. Koren, S., Walenz, B.P., Berlin, K., Miller, J.R., Bergman, N.H., Phillippy, A.M.: Canu: scalable and accurate long-read assembly via adaptive k-mer weighting and repeat separation. Genome research 27(5), 722–736 (2017)

13. Koslicki, D., Falush, D.: Metapalette: a k-mer painting approach for metagenomic taxonomic profiling and quantification of novel strain variation. MSystems 1(3) (2016)

14. Koslicki, D., Zabeti, H.: Improving minhash via the containment index with applications to metagenomic analysis. Applied Mathematics and Computation 354, 206–215 (2019)

15. LaPierre, N., Alser, M., Eskin, E., Koslicki, D., Mangul, S.: Metalign: efficient alignment-based metagenomic profiling via containment min hash. Genome biology 21(1), 1–15 (2020)

16. Li, P., König, A.C.: Theory and applications of b-bit minwise hashing. Communications of the ACM 54(8), 101–109 (2011)

17. Liu, B., Yuan, J., Yiu, S.M., Li, Z., Xie, Y., Chen, Y., Shi, Y., Zhang, H., Li, Y., Lam, T.W., et al.: Cope: an accurate k-mer-based pair-end reads connection tool to facilitate genome assembly. Bioinformatics 28(22), 2870–2874 (2012)

18. Luo, R., Liu, B., Xie, Y., Li, Z., Huang, W., Yuan, J., He, G., Chen, Y., Pan, Q., Liu, Y., et al.: Soapdenovo2: an empirically improved memory-efficient short-read de novo assembler. Gigascience 1(1), 2047–217X (2012)

19. Ondov, B.D., Starrett, G.J., Sappington, A., Kostic, A., Koren, S., Buck, C.B., Phillippy, A.M.: Mash screen: high-throughput sequence containment estimation for genome discovery. Genome biology 20(1), 1–13 (2019)

20. Ondov, B.D., Treangen, T.J., Melsted, P., Mallonee, A.B., Bergman, N.H., Koren, S., Phillippy, A.M.: Mash: fast genome and metagenome distance estimation using minhash. Genome biology 17(1), 1–14 (2016)

21. Pierce, N.T., Irber, L., Reiter, T., Brooks, P., Brown, C.T.: Large-scale sequence comparisons with sourmash. F1000Research 8 (2019)

22. Rana, S.B., Zadlock IV, F.J., Zhang, Z., Murphy, W.R., Bentivegna, C.S.: Comparison of de novo transcriptome assemblers and k-mer strategies using the killifish, fundulus heteroclitus. PLoS One 11(4), e0153104 (2016)

23. Sarmashghi, S., Bohmann, K., Gilbert, M.T.P., Bafna, V., Mirarab, S.: Skmer: assembly-free and alignment-free sample identification using genome skims. Genome biology 20(1), 1–20 (2019)

24. Schulz, M.H., Weese, D., Holtgrewe, M., Dimitrova, V., Niu, S., Reinert, K., Richard, H.: Fiona: a parallel and automatic strategy for read error correction. Bioinformatics 30(17), i356–i363 (2014)

25. Solomon, B., Kingsford, C.: Improved search of large transcriptomic sequencing databases using split sequence bloom trees. In: International Conference on Research in Computational Molecular Biology. pp. 257–271. Springer (2017)

26. Wood, D.E., Salzberg, S.L.: Kraken: ultrafast metagenomic sequence classification using exact alignments. Genome biology 15(3), 1–12 (2014)

27. Zhang, H., Zhu, L., Huang, D.S.: Wsmd: weakly-supervised motif discovery in transcription factor chip-seq data. Scientific reports 7(1), 1–12 (2017)

28. Zhang, Q., Jun, S.R., Leuze, M., Ussery, D., Nookaew, I.: Viral phylogenomics using an alignment-free method: A three-step approach to determine optimal length of k-mer. Scientific reports 7(1), 1–13 (2017)

